# BioOS: A Gene-Driven Digital Twin Runtime for Emergent Plant Development

**DOI:** 10.64898/2026.03.14.711542

**Authors:** Emeline Auger, Mickaël Gandecki, Charly Delarche, François-Xavier Heng

## Abstract

Predicting plant mutant phenotypes requires models that connect gene regulation to organ-scale morphogenesis without collapsing mechanism into phenomenological rules. We present **BioOS**, a curated mechanistic runtime built around the **Formal Cell**, a minimal signal-processing abstraction in which promoter evaluation, transcription, translation, and protein state drive cell division, differentiation, and elongation. A multi-scale architecture combining TissueUnits and FormalCells enables real-time simulation of bounded *Arabidopsis thaliana* developmental programs while keeping the primary claim anchored in primary-root auxin transport. On the official five-case root-auxin benchmark, BioOS achieves a 75.4% mean score, 5/5 qualitative matches, 5/5 quantitative passes, and Spearman severity correlation ρ = 0.70. The deployed auxin slice uses a curated 35-gene registry; for readability, this manuscript details an 18-gene core subnetwork. Beyond the primary auxin claim, the same runtime closes official cytokinin (5/5), flowering (5/5), and photosynthesis (7/7) gates, while a candidate root-patterning panel passes 8/8. BioOS should therefore be read as a benchmark-validated runtime for bounded developmental prediction rather than as a single-slice demonstration.

Key Resulty Results
The following results summarize the current validation status of BioOS on the primary-root auxin slice and its surrounding benchmark framework:

- 35-gene root-auxin runtime, with an 18-gene core GRN illustrated here
- Emergent division, differentiation, elongation from gene expression
- Official auxin panel closed: 5/5 qualitative matches, 5/5 full passes, 75.4% mean score, ρ = 0.70 severity ranking
- Four official gates closed in one runtime: root_auxin 5/5, cytokinin 5/5, flowering 5/5, photosynthesis 7/7
- Broader benchmark corpus: 6 suites / 63 cases; the 8-case candidate root-patterning panel currently passes 8/8
- Real-time capable: 175 TissueUnits + 200 cells = 8 ms/tick

## 1 Introduction

Predicting the phenotype of plant mutants from mechanistic first principles is a central goal of systems biology. Current approaches span coarse statistical models [Péret et al., 2012], detailed biochemical simulations [Band et al., 2012], and cell-based morphogenesis frameworks [Merks et al., 2011]. However, few offer both mechanistic traceability from genotype to phenotype *and* emergent prediction at organ scale.

The challenge is multi-scale: gene regulation operates at the molecular level (minutes, nanometers), while morphogenesis unfolds at the organ level (days, millimeters). Bridging these scales requires abstractions that preserve the essential regulatory logic without simulating every molecule. Most existing tools either operate at the biochemical level—making whole-organ simulation computationally intractable—or use phenomenological rules that bypass the genotype entirely.

We propose the **Formal Cell**: a radical simplification of the plant cell analogous to the formal neuron [McCulloch and Pitts, 1943]. Just as McCulloch and Pitts showed that simple threshold units could compute logical functions when connected in networks, we show that minimal gene-expression units can reproduce auxin-mediated root phenotypes—and that division, differentiation, and elongation *emerge* from the execution of a gene regulatory network rather than from explicit rules. The key insight is that the transfer function of the Formal Cell is gene expression itself: inputs (genome, environment, neighbor signals, current protein state) are processed by evaluating promoters and translating proteins, and the resulting protein concentrations determine all downstream behavior.

### 1.1 Contributions

This work makes the following contributions toward gene-driven, emergent plant simulation:

1. A **Formal Cell** abstraction with explicit gene expression runtime (promoter → mRNA → protein → behavior).
2. A **multi-scale architecture** with LOD switching (TissueUnit / FormalCell), enabling real-time simulation at 120 fps.
3. A **data-driven gene registry** (35 genes in the current root-auxin runtime; 18-gene core subset illustrated here) with promoter weights, TF bindings, kinetics, and epigenetic configuration.
4. **Emergent behavior**: division from cyclin/CDK, differentiation from PLT/KRP ratios, elongation from auxin response—no hardcoded thresholds.
5. A **benchmark-safe causal decoupling** of developmental zone labels, moving them from computation-time priors to post-hoc diagnostics while preserving the v1 causal core.
6. An **epigenetic memory model** for irreversible cell fate commitment.
7. A **three-level, multi-panel validation framework** spanning a six-suite, 63-case benchmark corpus with closed official gates in root auxin, cytokinin, flowering, and photosynthesis.
8. A **mechanistic extension path** for candidate panels such as root patterning, plasmodesmata, and intracellular auxin compartment chemistry without altering the causal core used by the official gates.

This framing matters for interpretation. BioOS is no longer best read as an early proof-of-concept around a single root demo. The primary publication claim remains the closed root-auxin gate because it most directly exercises the Formal Cell transport-and-growth core, but the surrounding runtime also closes official cytokinin, flowering, and photosynthesis slices. The paper therefore centers the auxin slice as the causal anchor while reporting the broader validation footprint explicitly.

## 2 The Formal Cell

### 2.1 Analogy with the Formal Neuron

The formal neuron computes *y* = *f*(Σ_*i*_ *w*_*i*_*x*_*i*_ + *b*): a weighted sum of inputs passed through an activation function. It is a radical simplification of a biological neuron, yet networks of formal neurons can learn complex functions. We adopt the same philosophy for plant cells.

The Formal Cell computes:

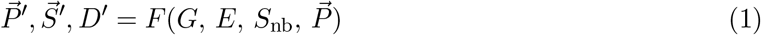

where *G* is the genome (immutable program), *E*(*t*) the local environment (auxin concentration, light, cytokinin, etc.), *S*_nb_(*t*) signals received from neighboring cells, and *P*(*t*) the current protein inventory. Outputs are *P*(*t*+1) (updated protein concentrations), *S*(*t*+1) (outgoing signals emitted to neighbors), and *D*(*t*+1) (division decision). The transfer function *F* is *entirely* gene expression: it evaluates promoters, produces mRNA, translates proteins, applies decay, and derives the three output quantities from the resulting protein state. No behavioral rule exists outside this pipeline.

### 2.2 Formal Cell vs. Real Cell

The Formal Cell is not a biophysical model—it is a *computational* model that retains exactly the regulatory logic required for emergent morphogenesis and discards the rest. Table 1 summarizes each simplification and the biological justification for accepting it. The central argument is that developmental decisions (division, fate, elongation) are determined by a small regulatory subnetwork, not by the full complement of ∼30 000 genes or by the physical details of organelles.

**Table 1.** Formal Cell simplifications relative to a real plant cell. In each case, the simplification is justified because the dropped detail is not necessary for emergent developmental regulation in the simulated domain.

| Real cell | Formal Cell | Why acceptable |
| --- | --- | --- |
| $\sim 30k$ genes | 35 curated runtime genes | Curated regulatory subnet drives development |
| Organelles | Protein type enum | Behavior captured by protein effects |
| 3D membrane | Abstract membrane | PIN polarity modeled explicitly |
| Continuous | Discrete tick ($\Delta t$) | Morphogenesis timescale: hours–days |
| $10^6+$ mol. | Concentration floats | Population-level sufficient |
| Complex cascades | 18-node core GRN / 35-gene runtime | Key motifs preserved |

### 3 Gene Expression Runtime

The core innovation of BioOS is that the Formal Cell’s transfer function is implemented as a *gene expression runtime*: a general-purpose engine that evaluates promoters, transcribes mRNA, translates proteins, and applies decay—for any set of genes defined in the registry. Unlike simulators that hardcode pathway logic in procedural code, BioOS computes the same pipeline for every gene at every tick, and behavioral differences between genotypes arise entirely from differences in the registry configuration.

### 3.1 Gene Expression Function

The expression rate of gene *g*_*i*_ at time *t* depends on the current transcription factor concentrations, the local environment, and a per-gene epigenetic accessibility factor. Specifically:

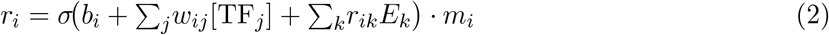

where σ is a sigmoid (1/(1 +*e*^*−x*^)), *b*_*i*_ the basal rate, *w*_*ij*_ the weight of transcription factor *j* on gene *i*(positive for activation, negative for repression), *r*_*ik*_ the sensitivity of the promoter to environmental signal *k*, and *m*_*i*_ ∈ [0, 1] the epigenetic accessibility factor (Section 8). This formulation follows the linear-additive thermodynamic model of transcriptional regulation [Bintu et al., 2005] with sigmoid saturation [Alon, 2019].

### 3.2 Transcription–Translation Cycle

Once the expression rate *r*_*i*_ is computed, mRNA and protein concentrations are updated each tick (Δ*t* in hours) via first-order kinetics. The decay terms are derived from experimentally measured half-lives, ensuring that mRNA and protein turnover rates are biologically grounded:

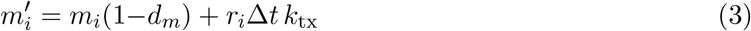

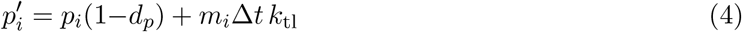

where *m*_*i*_ = [mRNA_*i*_], *p*_*i*_ = [Prot_*i*_], primes denote the state at 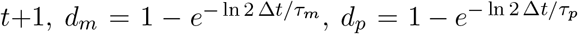, *k*_tx_ = 0.35, *k*_tl_ = 0.18.

The key emergent property of this architecture is that **proteins produced by one gene immediately become available as transcription factors for other genes** at the next tick. This creates closed regulatory loops such as ARF7→ PIN1→ auxin flux→ TIR1 → ARF7, where no single step is prescribed but the overall behavior emerges from iterated application of Equations 3–4.

## 4 Multi-Scale Architecture

A real *Arabidopsis* plant has approximately 10^8^ cells. Simulating each one individually with the full gene expression pipeline is computationally infeasible. BioOS addresses this with a multi-scale architecture in which the plant is represented as a directed tree of *segments*, and each segment can operate at one of two levels of detail (LOD) depending on its biological role.

At **LOD 0**, a segment is represented as a set of *TissueUnits*: coarse compartments corresponding to radial tissue layers. Each TissueUnit tracks molecular concentrations and integrates them forward in time using ordinary differential equations (ODEs). This mode is computationally efficient and covers the majority of the plant body (vascular tissue, elongation zone).

At **LOD 1**, a segment uses individual *FormalCells*: each cell runs the full gene expression pipeline (promoter evaluation, transcription, translation) every tick. This mode is reserved for meristems and user-inspected zones where cellular resolution is necessary. LOD switching is transparent and can be triggered dynamically during a simulation run.

### 4.1 Radial Symmetry Factorization

Stems and roots exhibit approximate rotational symmetry. Rather than simulating the full 3D cylinder (approximately 50 000 cells per cross-section), BioOS exploits this symmetry by simulating a single radial slice per segment. The slice consists of five concentric tissue layers—epidermis, cortex, endodermis, phloem, and xylem—each corresponding to one TissueUnit. The root is additionally split into upper and lower hemispheres to enable gravitropic asymmetry modeling, giving 10 TissueUnits per segment.

## 5 Gene Registry

The gene registry is the “program” that every Formal Cell executes. Rather than encoding pathway logic in simulator code, BioOS externalizes it: each gene is described in a standalone JSON file that specifies the promoter weights (which transcription factors activate or repress the gene, and by how much), kinetic parameters (mRNA and protein half-lives, transcription and translation efficiencies), the product protein type, and an optional epigenetic configuration. Adding a new gene to the simulation requires only a new JSON file; no Rust code needs to change.

The current root-auxin runtime contains 35 curated genes. For readability, Table 2 lists an 18-gene core subset spanning four functional families: auxin transport and response (9 genes), the extended auxin signaling network (3 genes), the cell cycle (5 genes), and meristem identity (1 gene). This illustrated core is the regulatory subnetwork used throughout the manuscript figures and explanations, while the deployed runtime also includes additional regulators, transport modifiers, and benchmark extension proxies.

**Table 2.** Illustrated core gene registry (18 of 35 current root-auxin genes). Product protein, mRNA/protein half-lives (τ), TF inputs (+ = activation, − = repression), and environmental signal inputs.

| Gene | Product | $\tau_m$ (h) | $\tau_p$ (h) | TF inputs | Env inputs |
| --- | --- | --- | --- | --- | --- |
| *Auxin transport & response (9 genes)* |  |  |  |  |  |
| AUX1 | Aux1 influx | 2.0 | 8.0 | ARF7 (+) | Auxin |
| PIN1 | Pin1 efflux | 2.5 | 8.5 | ARF7 (+) | Response |
| PIN2 | Pin2 efflux | 2.5 | 8.5 | ARF7 (+) | Auxin |
| PIN3 | Pin3 lateral | 3.0 | 10.0 | — | Gravity, Auxin |
| TAA1 | Taa1 biosyn. | 2.0 | 7.5 | — | Auxin, Nutrient |
| YUC1 | Yuc1 biosyn. | 2.5 | 8.5 | — | Auxin |
| TIR1 | Tir1 receptor | 3.0 | 10.0 | — | Auxin |
| AXR1 | Axr1 signal. | 3.0 | 10.0 | TIR1 (+) | Response |
| ARF7 | Arf7 TF | 2.5 | 7.5 | TIR1 (+) | Auxin, Response |
| *Extended auxin network (3 genes)* |  |  |  |  |  |
| IAA17 | IAA17 repr. | 1.5 | 4.0 | ARF7 (–) | Auxin |
| AFB2 | AFB2 receptor | 3.0 | 9.0 | — | Auxin |
| ARF19 | ARF19 TF | 2.5 | 7.5 | TIR1 (+) | Auxin, Response |
| *Cell cycle (5 genes)* |  |  |  |  |  |
| CDKA | Cyclin A | 2.0 | 6.0 | E2F (+) | Auxin |
| CDKB | Cyclin B | 2.0 | 5.0 | E2F (+) | Auxin |
| KRP | CDK inhibitor | 2.5 | 7.0 | — | Auxin (–) |
| RBR | Retinoblastoma | 3.0 | 8.0 | — | — |
| E2F | E2F TF | 2.0 | 5.0 | — | Auxin |
| *Meristem identity (1 gene)* |  |  |  |  |  |
| PLT | Plethora TF | 3.0 | 9.0 | ARF7 (+) | Auxin |

Adding a new gene requires only a single JSON descriptor. The following example shows the complete PIN1 specification: it defines the product type, half-lives, and promoter wiring (ARF7 as activating TF, auxin response as environmental input):

~~~
{ “gene_id ”: “PIN1 “, “product ”: “PIN1 “,
  “basal_rate ”: 0.08,
  “mrna_half_life_hours ”: 2.5,
  “protein_half_life_hours ”: 8.5,
  “promoter ”: {
    “tf_bindings ”: [
     {“tf”:” ARF7 “, “weight ”:0.14}],
    “env_responses ”: [
     {“signal ”:” Response “, “coeff ”:0 .16 }]
} }
~~~

## 6 Gene Regulatory Network

A core 18-gene subset of the current 35-gene root-auxin runtime is shown in Figure 7. These genes do not operate in isolation—they form an interconnected regulatory network in which the protein produced by one gene modulates the expression of others. The network contains two types of regulatory edges: activation edges (solid arrows in Figure 7), in which the source protein increases promoter activity of the target gene, and repression edges (dashed arrows), in which the source protein decreases promoter activity. The topology can be organized into four modules: auxin signaling (TIR1, AFB2, AXR1, ARF7, ARF19, IAA17), auxin transport (PIN1, PIN2, PIN3, AUX1),auxin biosynthesis (TAA1, YUC1), and cell cycle/identity (CDKA, CDKB, KRP, RBR, E2F, PLT). These modules are interconnected: for example, ARF7 (signaling module) activates PIN1 and PIN2 (transport module), and PLT (identity module) is activated by ARF7 and maintained by auxin. It is this cross-module wiring that gives rise to emergent phenotypes: a single gene knockout disrupts multiple modules simultaneously, producing the complex developmental defects seen in auxin mutants.

**Figure 1.**
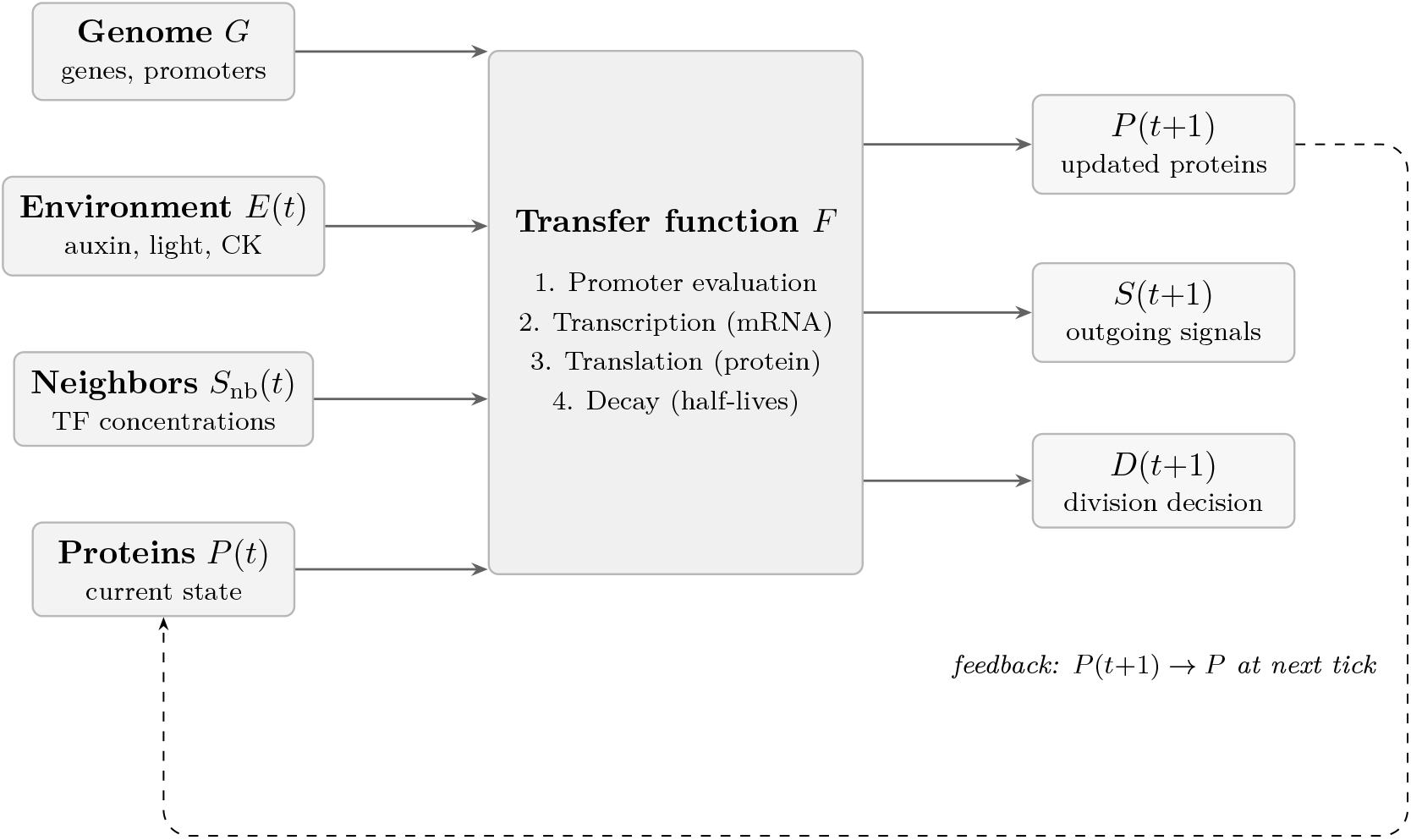
The Formal Cell. Four inputs (genome, environment, neighbor signals, current protein state) feed the transfer function *F*, which evaluates gene promoters, produces mRNA, translates proteins, and applies decay half-lives. Three outputs are produced each tick: updated proteins, outgoing signals, and a division decision. The dashed feedback arrow indicates that the protein state produced at tick *t* becomes an input at tick *t*+1, enabling dynamic regulatory circuits such as the TIR1–ARF7–PIN1 feedback loop.

**Figure 2.**
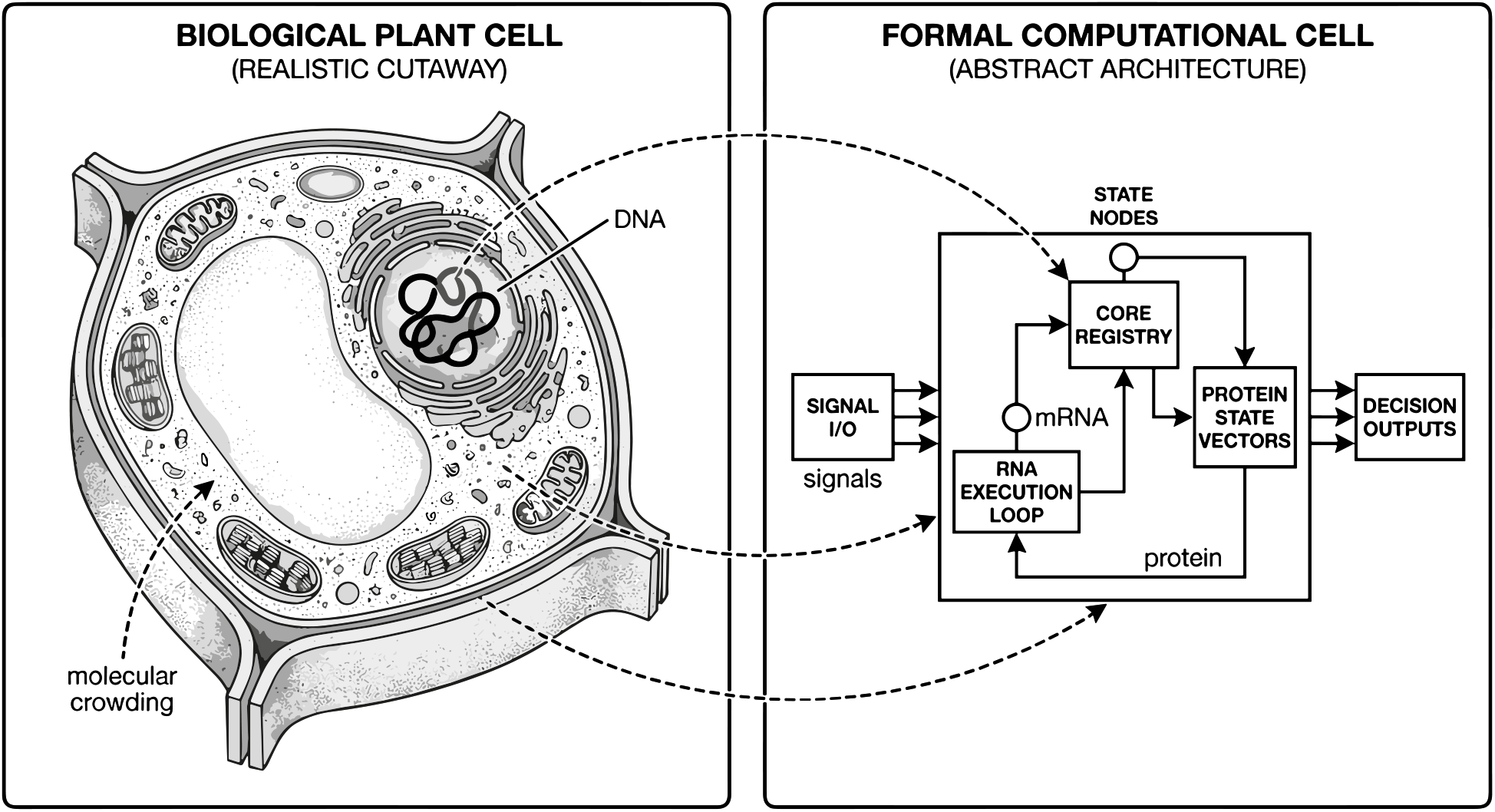
Biological plant cell versus formal computational cell. The left panel shows biophysical complexity (organelles, membrane, crowding), while the right panel shows the executable abstraction used by BioOS (RNA execution loop, protein state vectors, signal I/O, and decision outputs). The mapping highlights what is simplified and what is preserved for mechanistic emergence.

**Figure 3.**
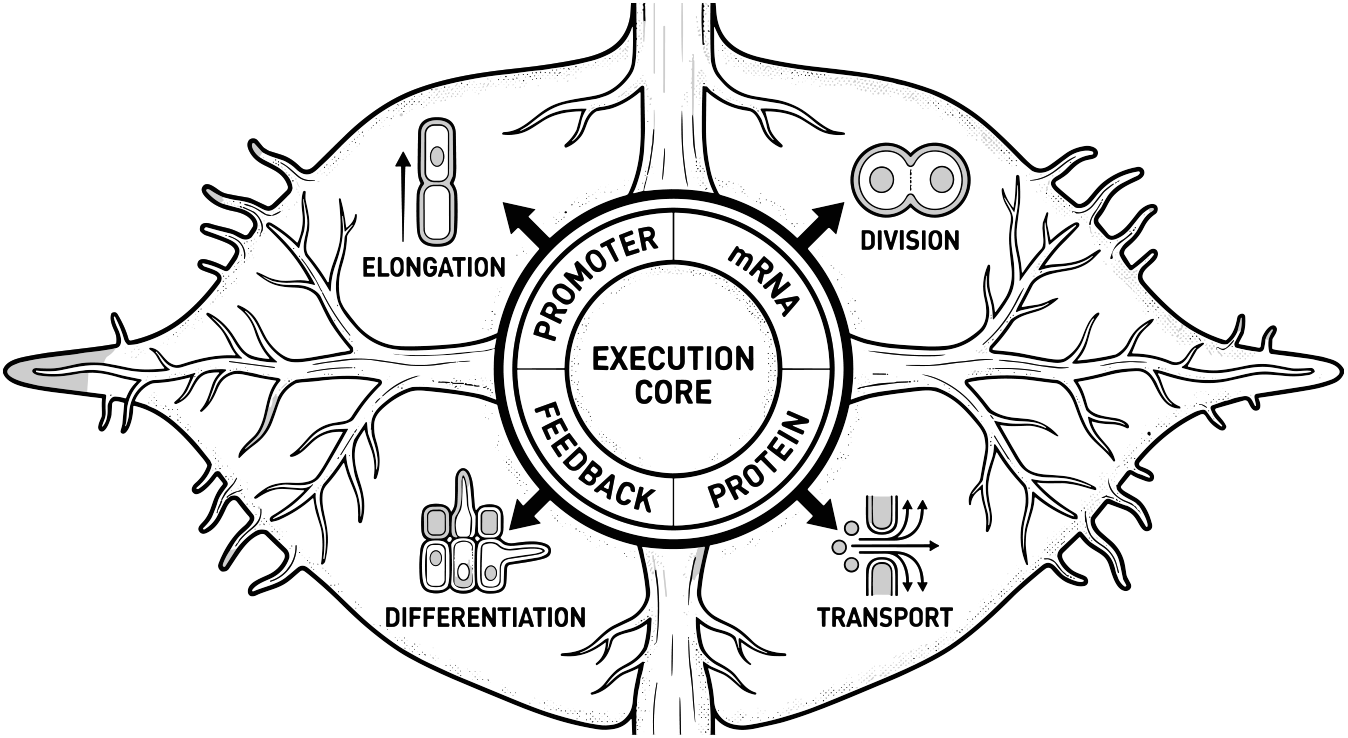
RNA execution core inside the Formal Cell. Promoter evaluation, transcription, translation, and feedback form a recurrent computational engine that drives transport, division, elongation, and differentiation outputs.

**Figure 4.**
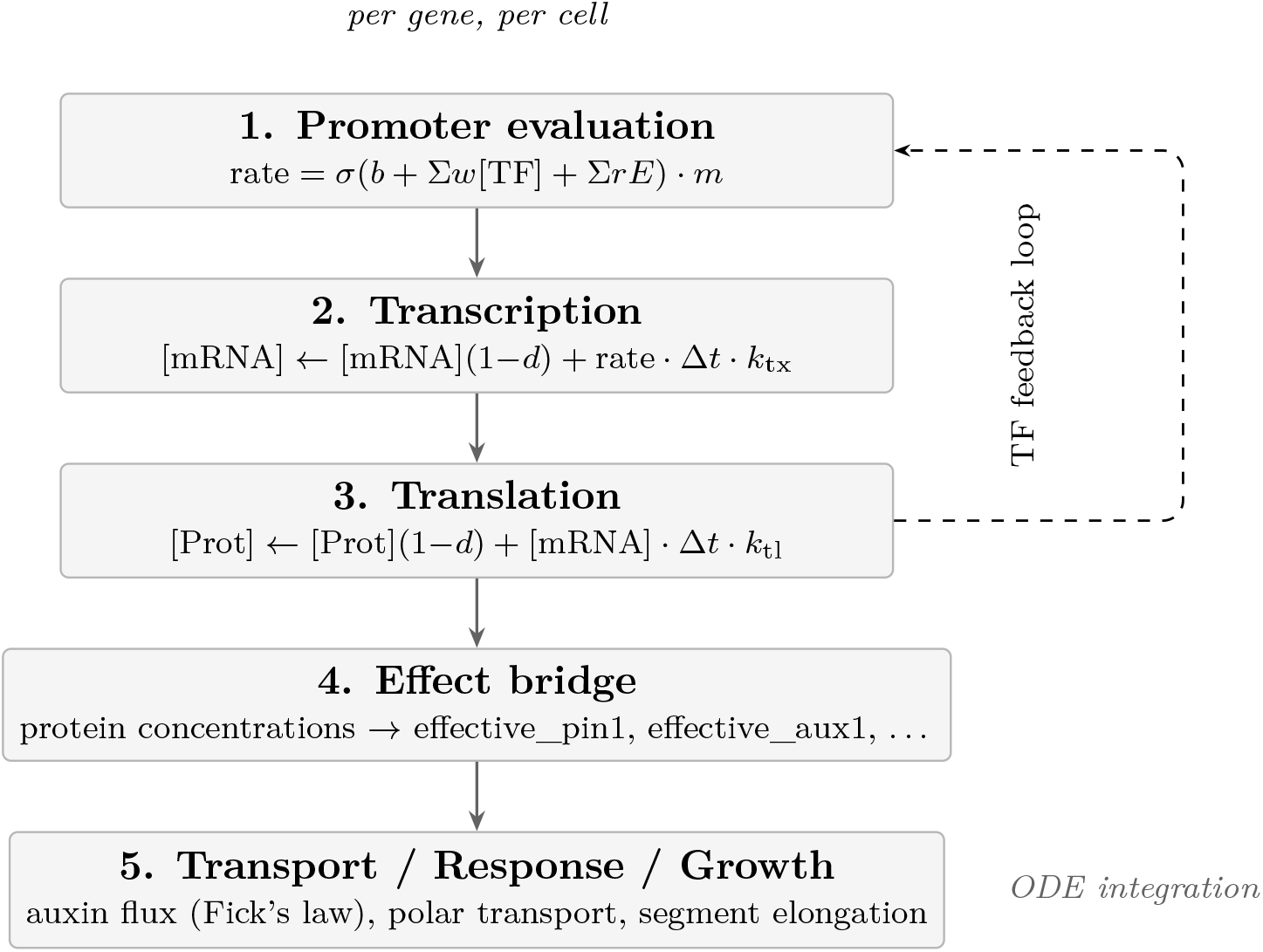
Gene expression pipeline. Five stages execute per cell per tick. Stages 1–3 operate once per gene: the promoter is evaluated, mRNA is updated, and protein concentration is updated. Stage 4 maps protein concentrations to functional effectors (e.g. PIN1 concentration becomes the polar auxin export rate). Stage 5 integrates transport and growth. The dashed arrow represents the TF feedback loop: proteins produced in stage 3 re-enter stage 1 at the next tick, creating dynamic regulatory circuits without any separate wiring code.

**Figure 5.**
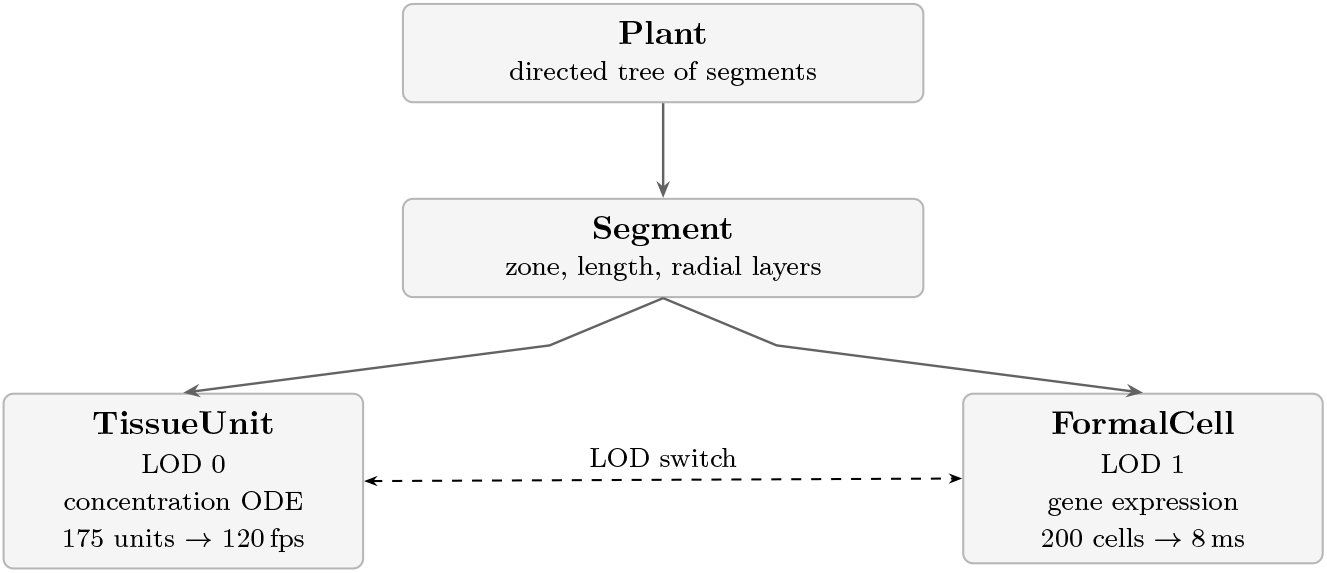
Multi-scale hierarchy. The plant is a tree of segments. Each segment operates at LOD 0 (TissueUnit, concentration-based ODE) or LOD 1 (FormalCell, full gene expression runtime). Meristems run at LOD 1; bulk tissue runs at LOD 0. The bidirectional arrow indicates that LOD switching is dynamic and transparent to the rest of the simulation. Combined, 175 TissueUnits and 200 meristem cells yield 8 ms per tick—sufficient for 120 fps in WASM.

**Figure 6.**
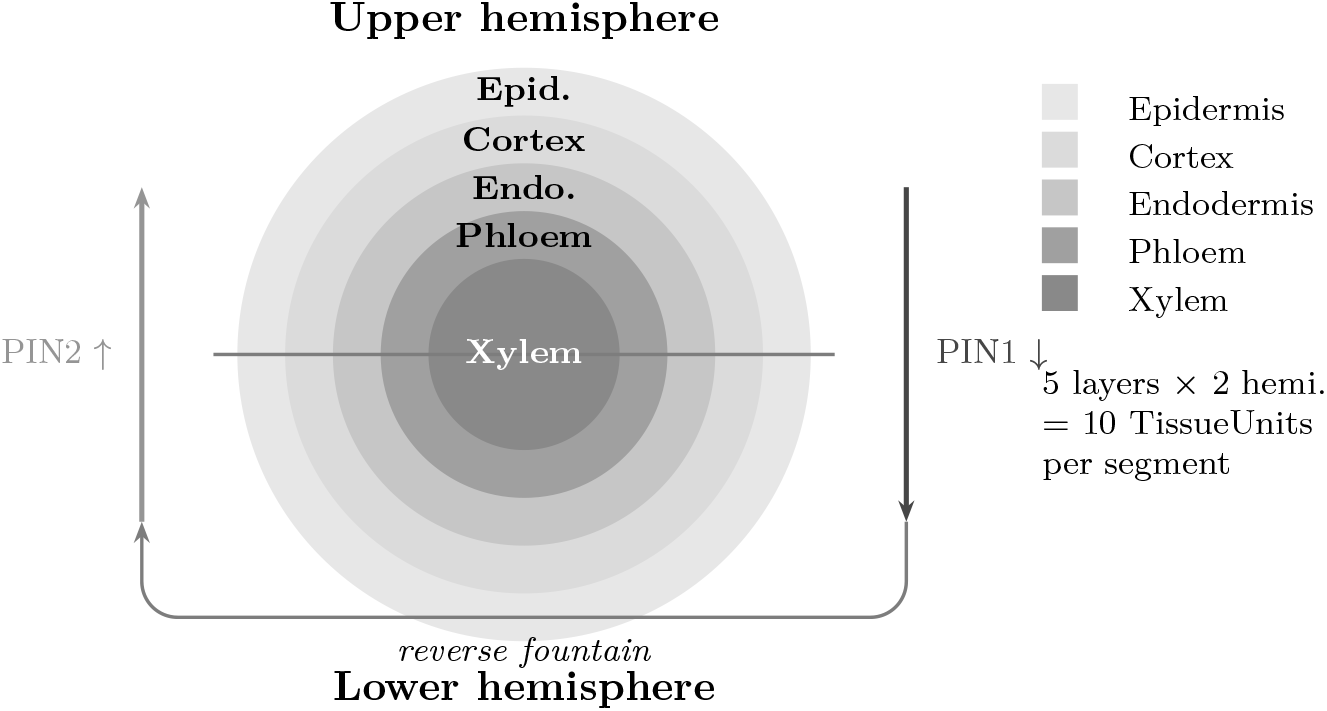
Radial cross-section of a root segment showing the five concentric tissue layers. Gray shading distinguishes the layers: the outermost epidermis is lightest; the xylem core is darkest. Auxin flows rootward through the stele via PIN1 efflux carriers (solid arrow, right) and shootward through the epidermis via PIN2 (lighter arrow, left), creating the reverse fountain pattern [Grieneisen et al., 2007]. The horizontal dividing line separates upper and lower hemispheres, enabling gravitropic bending to be modeled by asymmetric auxin distribution across the two halves.

**Figure 7.**
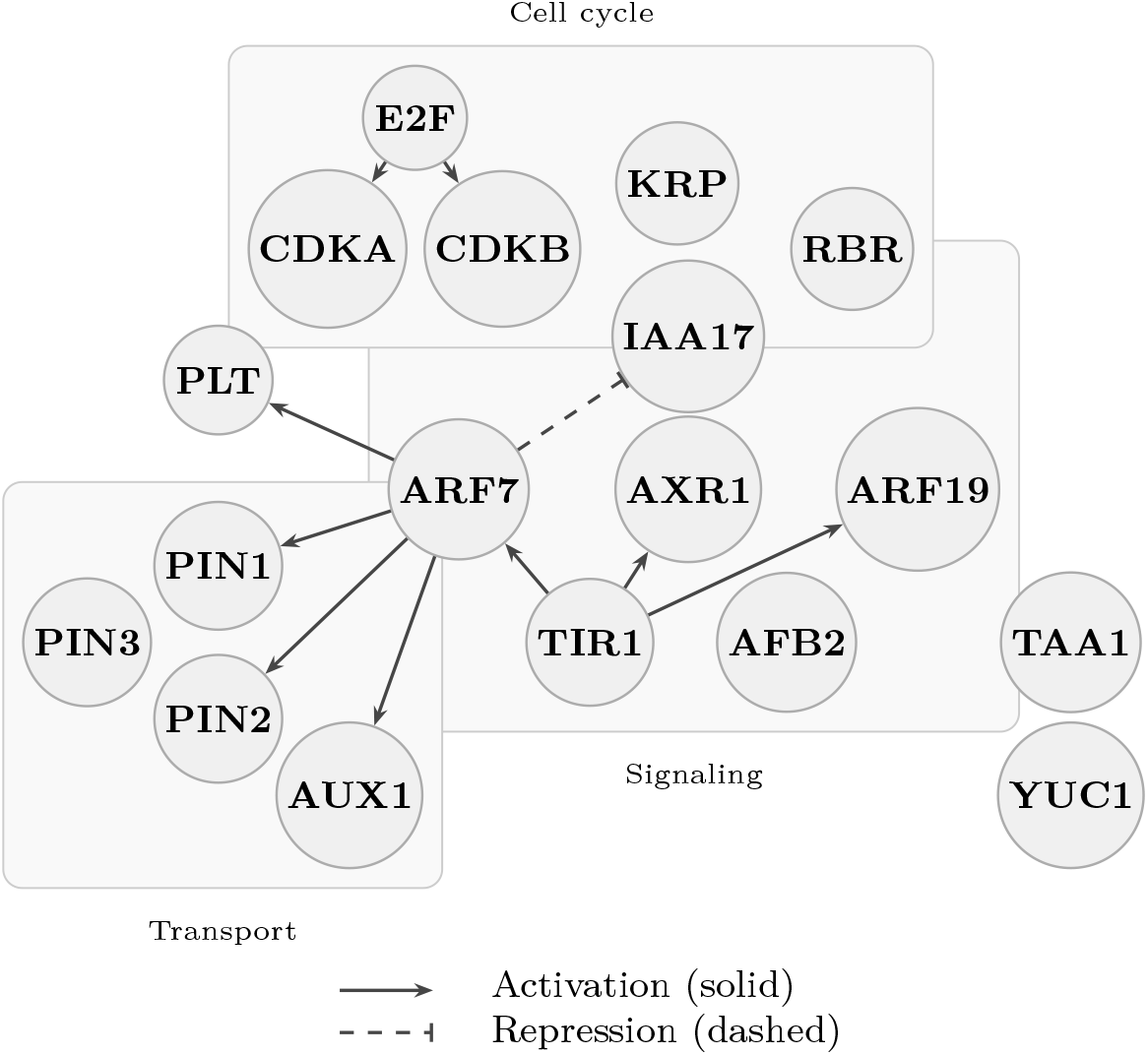
Core gene regulatory network topology (18 highlighted genes from the current 35-gene root-auxin runtime). Solid arrows indicate activation (the source protein increases expression of the target gene); dashed arrows indicate repression. Background rectangles group genes into functional modules: signaling (center), transport (left), and cell cycle (top). Biosynthesis genes TAA1 and YUC1 (right) and meristem identity gene PLT (upper left) are not enclosed in module boxes for visual clarity. The network structure is what enables emergent phenotypes: disrupting any node affects all its downstream targets without any rule being explicitly changed.

**Figure 8.**
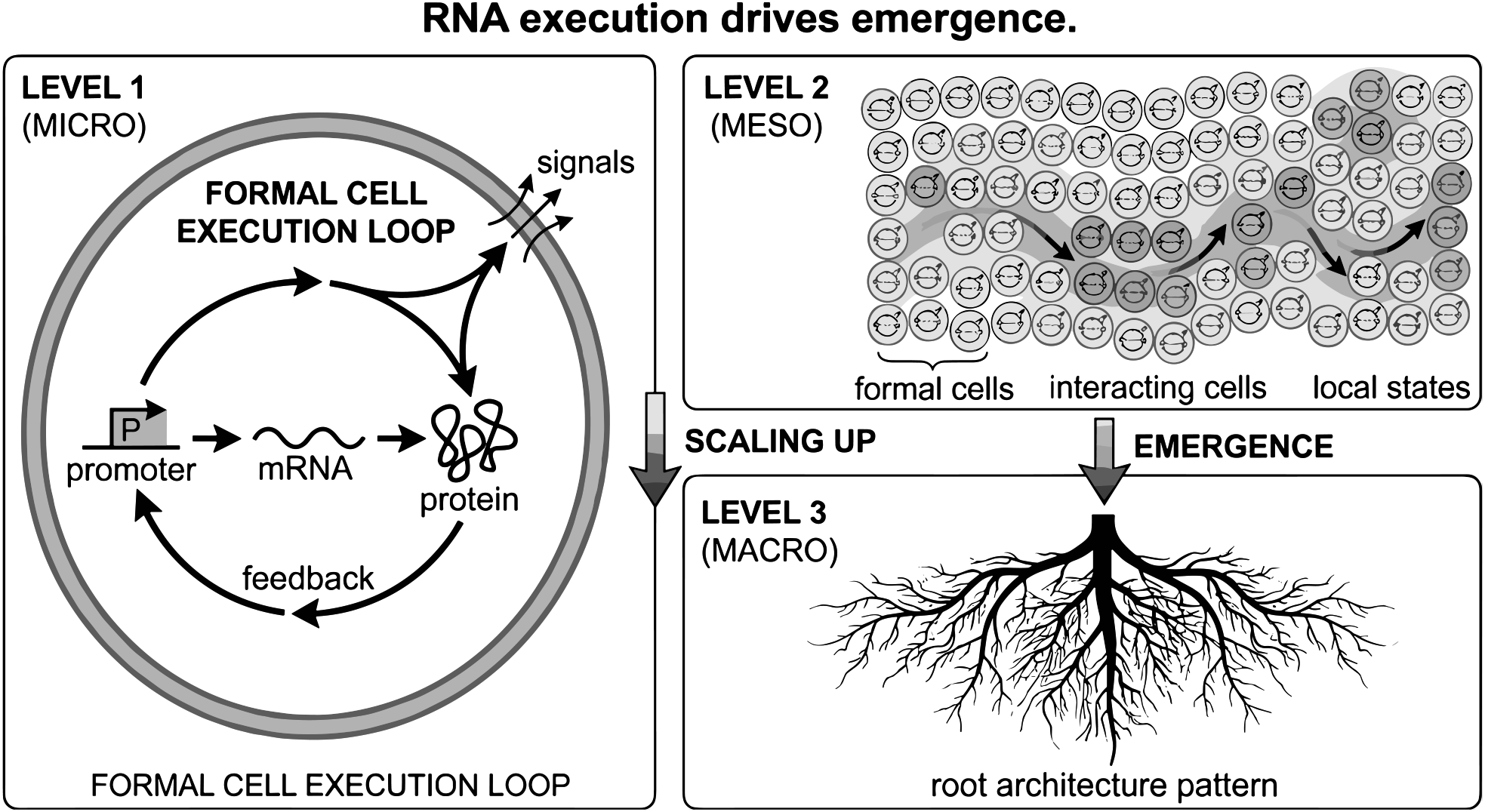
Emergence across scales from the same RNA execution rule. At the micro level, each Formal Cell runs the promoter→ mRNA →protein loop; at the meso level, interacting cells form local state patterns; at the macro level, root-scale architecture emerges as a collective consequence rather than a hardcoded template.

## 7 Emergent Behavior

Cell behavior *emerges* from gene expression rather than being prescribed. This section describes how the three primary developmental behaviors—division, differentiation, and elongation—arise as consequences of the protein concentrations computed by the gene expression runtime, with no dedicated behavioral rules in the simulator.

### Design Principle

Emergence comes from the Formal Cell itself. Division, differentiation, and elongation arise from protein concentrations computed by the gene expression runtime. A knockout mutant shows an altered phenotype because its protein is absent from the regulatory network—not because a rule was disabled. The genotype alone determines the phenotype, through the execution of the gene expression pipeline.

### 7.1 Emergent Division

Cell division is controlled by five cell cycle genes (CDKA, CDKB, KRP, RBR, E2F), whose interactions form a well-characterized bistable switch in *Arabidopsis* [De Veylder et al., 2007]. The mechanism operates as follows:

- **CDKA/CDKB** (cyclins): both genes are activated by E2F and further induced when auxin is present. Their protein products act as the “gas pedal” of the cell cycle: high CDK activity drives S-phase entry.
- **KRP** (CDK inhibitor): this gene is repressed by auxin, so KRP protein accumulates when auxin drops, acting as the “brake” that prevents division in differentiated cells.
- **RBR**: sequesters the E2F transcription factor in its unphosphorylated state. When CDK activity is sufficient to phosphorylate RBR, E2F is released and further activates CDKA and CDKB, forming a positive feedback loop.
- **Division**: occurs when the integrated CDK signal exceeds the KRP brake—no hardcoded threshold exists in the code; the threshold emerges from the balance of these concentrations.

### 7.2 Emergent Differentiation

Cell fate is determined by transcription factor ratios rather than by explicit state transitions. The key player is PLT (Plethora), which maintains meristematic identity [Aida et al., 2004]. When auxin concentration is high (near the root tip), ARF7 activates PLT expression, keeping cells undifferentiated and division-competent. As cells move away from the tip and local auxin drops, ARF7 activity falls, PLT expression decreases, and differentiation begins. Because the transition arises from a continuous change in TF ratios rather than a discrete rule, there is no fixed boundary between meristem and elongation zone—the gradient itself is an emergent property. The SHR/SCR radial patterning module, which specifies endodermis identity [Helariutta et al., 2000], also appears in the candidate root-patterning panel as a mechanistic extension, while the publication-default root-auxin gate continues to center on the PLT/auxin differentiation axis.

### 7.3 Emergent Elongation

Root elongation emerges from the auxin response chain: auxin activates TIR1, which in turn promotes AXR1 and ARF7, and ARF7 confers growth competence on the cell. Mutants that disrupt any step of this chain—*aux1* (reduced auxin influx), *axr1* (impaired signaling), or *pin2* (impaired efflux and thus altered auxin gradient)—show reduced elongation as an emergent consequence of the missing protein, without any dedicated elongation rule being disabled.

## 8 Epigenetic Memory

A purely concentration-based model has a fundamental limitation with respect to cell identity: because protein concentrations are continuous and reversible, a cell that has differentiated could in principle re-dedifferentiate if its protein environment changed. In biology, however, differentiation is largely irreversible due to epigenetic modifications—in particular, Polycomb-mediated H3K27me3 methylation, which silences developmental genes by compacting chromatin regardless of the transcription factor environment [Angel et al., 2011].

BioOS models epigenetic state through a per-gene, per-cell accessibility factor *m*(*g*_*i*_) ∈ [0, 1] that scales the expression rate in Equation 2:

- *m* = 1: the promoter is fully open; the gene responds normally to transcription factor and environmental signals.
- *m* = 0: the promoter is silenced; the gene is unresponsive regardless of activating signals— modeling constitutive chromatin compaction.
- **Commitment**: when a differentiation-promoting TF (e.g. PLT) falls below a threshold for *N* consecutive ticks, antagonist genes are progressively silenced by decreasing *m* at a fixed rate. This is analogous to Polycomb-mediated H3K27me3 silencing and captures the irreversibility of fate commitment.

The epigenetic parameters are specified per gene in the registry:

~~~
“epigenetics ”: {
 “silencing_threshold ”: 0.2,
 “commitment_ticks ”: 48,
 “silencing_rate ”: 0.01,
 “reactivation_resistance ”: 0.95
}
~~~

These fields have the following meaning:

- `silencing_threshold:` TF level below which progressive silencing begins.
- `commitment_ticks:` number of consecutive ticks the threshold must be crossed before silencing starts (approximately 48 hours at a one-hour tick).
- `silencing_rate:` per-tick decrease in *m*, analogous to gradual chromatin compaction.
- `reactivation_resistance:` difficulty of reversing an established silenced state.

## 9 Validation

### 9.1 Primary Claim and Benchmark Footprint

The publication-default claim is anchored in a five-case primary-root auxin panel chosen to cover the core transport-and-response pathway at multiple levels: auxin influx (*aux1*), auxin efflux (*pin2/eir1*), auxin signaling (*axr1*), and auxin biosynthesis (*taa1/wei8*), plus the wild type (Col-0) as calibration reference. The panel is split into calibration cases (used to fit parameters), validation cases (held out during fitting and used to test generalization), and a holdout case (not shown to the system until final evaluation). This panel remains central because it exercises the full multi-scale Formal Cell transport-and-growth loop with a clean calibration/validation/holdout structure. Table 3 summarizes the five cases.

**Table 3.** Core benchmark panel: *Arabidopsis* primary root auxin mutants [Berardini et al., 2015]. The “Split” column indicates how each genotype is used: Calib. = calibration, Valid. = validation, Holdout = final evaluation.

| Genotype | Lesion | Growth | Split |
| --- | --- | --- | --- |
| Col-0 | Wild type | Normal | Calib. |
| *aux1* | Reduced influx | Reduced | Calib. |
| *pin2/eir1* | Impaired efflux | Reduced | Valid. |
| *axr1* | Impaired signaling | Reduced | Valid. |
| *taa1/wei8* | Reduced biosyn. | Reduced | Holdout |

This five-case panel remains the publication-default mechanistic claim because it is the slice most directly tied to the full Formal Cell causal core. It sits within a broader benchmark corpus of six suites and 63 cases. Outside `root_auxin`, official gates are also closed for `flowering (5/5), photosynthesis (7/7)`, and `cytokinin` (5/5, 74.3% mean score). The candidate eight-case `root_patterning` panel also passes 8/8 and exercises post-hoc zone diagnostics, persistent plasmodesmata (*PDBG1*), and intracellular auxin compartment perturbations (*WAT1, PILS5*).

### 9.2 Three-Level Validation

Validation is organized at three levels of mechanistic depth. Level 1 captures macroscopic phenotype (root length, growth class); Level 2 probes the underlying transport dynamics that generate the phenotype; Level 3 evaluates the temporal trajectory of the simulation over multiple days. This hierarchy ensures that correct phenotypes are not produced by incorrect mechanisms. Table 5 describes the metrics at each level.

### 9.3 Official Root-Auxin Results

The metrics in this subsection refer only to the official five-case primary-root auxin panel after calibration on Col-0 and *aux1*. They do not summarize the entire benchmark corpus; the broader cross-suite status is reported separately in Table 4. On this publication-default auxin slice, BioOS matches the expected qualitative growth/response class for all five genotypes, and all five pass the full quantitative gate. The tightest margins are concentrated in *axr1* and the *taa1/wei8* holdout, so this result should be read as a validated official slice with identifiable weak margins rather than as a finished calibration endpoint. The severity-ranking metrics remain useful as secondary diagnostics of mutant ordering, but the decisive criterion is closure of the per-case mechanistic gate itself.

**Table 4.** Current benchmark-gate status for the main slices discussed in the codebase.

| Suite | Scope | Status | Mean score |
| --- | --- | --- | --- |
| **root_auxin** | Official | 5/5 passed | 75.4% |
| **flowering** | Official | 5/5 passed | 93.2% |
| **photosynthesis** | Official | 7/7 passed | 86.2% |
| **cytokinin** | Official | 5/5 passed | 74.3% |
| **root_patterning** | Candidate | 8/8 passed | 86.8% |

**Table 5.** Validation levels with increasing mechanistic depth. A simulation must pass all three levels to be considered a mechanistically valid reproduction of a mutant phenotype.

| Lv. | Name | Metrics |
| --- | --- | --- |
| L1 | Phenotype | Root length ratio vs. Col-0, elongation rate, meristem size, qualitative growth class (normal / reduced / absent), qualitative auxin response class |
| L2 | Transport | Tip/base auxin gradient ratio, rootward and shootward flux magnitudes, reverse-fountain balance score |
| L3 | Trajectory | Day-by-day root length curve, auxin concentration time series, auxin response time series, growth competence trajectory |

#### Validation Results

- **Mean score**: 75.4% (weighted L1/L2/L3), mean coverage 97.3%
- **Qualitative**: 5/5 match expected growth & response class
- **Quantitative**: 5/5 pass all gates; 0/5 partial, 0/5 failing
- **Severity ranking**: Spearman *ρ* = 0.70, Kendall *τ* = 0.60

The remaining engineering gaps are no longer official-gate failures but margin management problems: *axr1* and *taa1/wei8* remain the most fragile passing cases, especially in transport- and trajectory-derived subscores, so they define the most informative frontier for future recalibration.

Taken together with Table 4, this means the paper gives a detailed mechanistic reading of the auxin slice while situating it inside a broader runtime that also closes three other official gates.

## 10 System Architecture

BioOS is implemented as a set of Rust crates, each with a single clearly bounded responsibility. The `kernel` crate is the simulation core; the remaining crates provide supporting capabilities. Table 6 summarizes the crate structure. All crates compile to both native Rust (for batch screening and testing) and WebAssembly (for the browser-based digital twin front end).

**Table 6.** BioOS crate structure (Rust 2021, stable toolchain).

| Crate | Responsibility |
| --- | --- |
| kernel | FormalCell, Nucleus, Gene, Segment, Plant, Scheduler; the multi-scale tick loop and LOD switching logic |
| biology | CuratedGenomeModel, gene registry loader, protein effect bridge (concentration to effector) |
| signals | Diffusion solver (Fick’s law), polar auxin transport (PIN model), xylem/phloem long-distance transport |
| environment | EnvironmentProfile, GROW LED program mapping, gravitropic vector |
| validation | Benchmark panel definitions, L1/L2/L3 phenotype scoring, Spearman severity correlation |
| renderer | Segment-level SVG/HTML diagnostic renderer, cross-section heatmaps, benchmark report generation |

## 11 Comparison with Existing Tools

Framework comparisons are easy to over-broaden. Here we narrow the scope to two plant-focused references that are closer to BioOS use cases: VirtualLeaf [Merks et al., 2011] and OpenAlea [Pradal et al., 2008]. This is a *positioning* table, not a leaderboard: each platform optimizes for a different scientific question.

VirtualLeaf is strong for cell mechanics and tissue growth rules; OpenAlea is strong for composable structural-functional pipelines and model integration. BioOS targets a different axis: executable gene-expression dynamics tied to a bounded mutant benchmark under real-time constraints. Table 7 summarizes these complementary emphases.

**Table 7.** Feature comparison with plant simulation frameworks. ✓ = supported, ✗ = not supported.

|  | BioOS | V.Leaf | OpenAlea |
| --- | --- | --- | --- |
| Gene expression runtime (promoter→mRNA→protein) | ✓ | ✗ | ✗ |
| Primary emphasis on cell mechanics | ✗ | ✓ | ✗ |
| Pipeline composability for structural-functional models | ✗ | ✗ | ✓ |
| Built-in mutant benchmark workflow (explicit pass/fail panel) | ✓ | ✗ | ✗ |
| Hybrid LOD for real-time interactive execution | ✓ | ✗ | ✓ |
| WASM/browser real-time target in core workflow | ✓ | ✗ | ✗ |

The key message is complementarity, not replacement: BioOS contributes a gene-execution-centric and benchmark-driven runtime, while existing frameworks remain valuable for mechanical detail or broader model orchestration.

## 12 Biological Debugger

One of the key intended applications of BioOS is **variant screening**: running hundreds or thousands of simulated genotypes and comparing their developmental trajectories to identify promising candidates for experimental validation. Standard phenotype-level comparison (“mutant A has shorter roots than mutant B”) does not explain *why* the phenotypes differ. The biological debugger addresses this by enabling mechanistic comparison at the level of individual gene executions.

The debugger concept introduces three types of breakpoints:

1. **Gene breakpoints**: the simulation pauses when a specified gene’s expression rate or protein concentration crosses a user-defined threshold in any cell. This allows the user to observe exactly when and where a particular protein becomes active.
2. **Cell breakpoints**: the simulation pauses when any cell undergoes a qualitative state transition (e.g. meristematic → elongating, or division → differentiation).
3. **Diff mode**: two genotypes are run in lockstep; the simulation auto-pauses at the first tick and cell where their protein states diverge beyond a user-defined tolerance.

Diff mode in particular transforms variant screening from a phenotype-level comparison to a *mechanistic* one: the user can identify the first molecular divergence event—which gene, in which cell, at which developmental stage—rather than only observing the phenotypic output.

## 13 Applications

BioOS supports four primary application domains, all built on the same underlying gene-driven runtime. Figure 9 illustrates these domains and their connection to the core runtime. The runtime’s data-driven design means that each application is enabled by configuration rather than by additional code: batch variant screening requires only a list of modified gene registry files; in-silico CRISPR prediction requires only a knockout or knockdown parameter on a specific gene.

**Figure 9.**
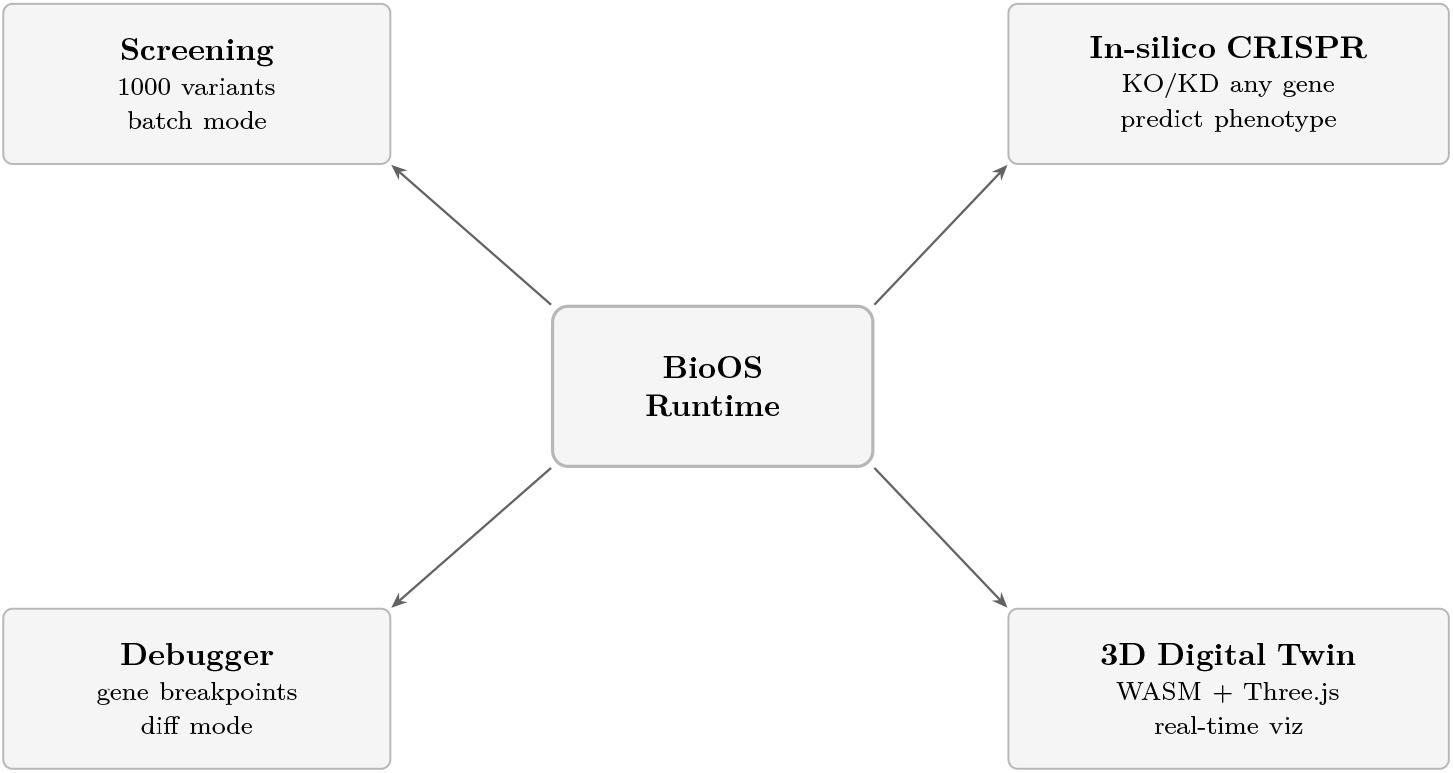
Application domains enabled by the BioOS gene-driven runtime. The central runtime connects to four use cases: high-throughput variant screening (batch mode, 1000 genotypes); in-silico CRISPR prediction (knockout or knockdown any registry gene); the biological debugger (gene-level breakpoints and diff mode for mechanistic comparison); and the 3D digital twin (real-time WebAssembly visualization via Three.js). All four modes use the same runtime and differ only in their input configuration.

## 14 Discussion

### 14.1 Strengths

The core strength of BioOS is *mechanistic traceability*: every phenotypic observation can be traced back to a specific sequence of gene expression events. The following properties characterize the current system:

- **Mechanistic traceability**: gene edit→ promoter→ expression→ protein→ transport → phenotype. Every step is observable and inspectable.
- **Validated breadth**: the same runtime closes official root-auxin, cytokinin, flowering, and photosynthesis gates, while the candidate root-patterning panel passes 8/8.
- **Architectural maturation**: the root runtime decouples developmental zone labels from the causal core, exporting computed zones as diagnostics rather than using them as next-tick inputs.
- **Mechanistic extensibility**: candidate root-patterning benchmarks exercise persistent plasmodesm permeability and intracellular auxin compartment summaries without reintroducing zone priors into the causal engine.
- **Developmental extension**: the engine has progressed beyond the static root slice toward seed-to-plant execution, with a causal germination GRN, state-driven seedling growth, and recursive lateral-root topology handled directly in the kernel.
- **Extensibility**: adding a gene, a pathway, or a tissue type is a data task rather than a code task.

### 14.2 Current Limitations

The following limitations are acknowledged and constitute the immediate development priorities:

- Cell cycle genes are defined in the registry but not yet fully calibrated against experimental division rate data; division emerges correctly in direction but not in magnitude.
- The closed root-auxin gate still has identifiable weak-margin cases: *axr1* and *taa1/wei8* are informative passes rather than failures, but they remain the most sensitive frontier for regression.
- The zone-decoupling refactor is complete and benchmark-safe at the engine level, but the new observational zone summaries are still best treated as diagnostic outputs rather than publication-default benchmark targets.
- Seed-to-plant execution, causal germination, and recursive lateral-root growth are present in the runtime, but they do not yet carry the same benchmark weight as the four currently closed official slices.
- Persistent plasmodesm connections and intracellular auxin compartment pools are implemented as candidate mechanistic extensions, but their experimental calibration remains thinner than the official auxin transport and signaling slice.
- Cytokinin, flowering, and photosynthesis close their official gates, but they remain more compact and less cell-mechanistic than the root-auxin kernel that anchors the main claim.
- Transcription is deterministic; stochastic noise in promoter firing is planned for the next runtime version.

### 14.3 Future Directions

- Convert benchmark closure into prospective biology by testing predicted perturbations *in vivo*, starting with coupled auxin-compartment mutants such as *wat1 pils5* and related epistasis panels.
- Increase the number of simulated cells and the scale of screening runs by vectorizing the scheduler while preserving the same causal semantics, debugger visibility, and benchmark traceability as the sequential path.
- Continue hardening the weakest-margin official cases—especially *axr1* and *taa1/wei8* in root auxin—so that in vivo prediction work rests on a stable benchmark base.

## 15 Conclusion

BioOS shows that a curated gene-expression runtime can reproduce bounded plant mutant phenotypes with mechanistic traceability and real-time execution. In the primary-root auxin slice, the Formal Cell couples promoter logic, transcription, translation, transport, and growth strongly enough to close the official five-case benchmark, with 5/5 qualitative matches, 5/5 quantitative passes, and a 75.4% mean score.

That core claim sits inside a broader validated runtime. The same engine also closes official cytokinin, flowering, and photosynthesis gates, while candidate root-patterning cases extend the mechanism to post-hoc zones, plasmodesmata, and intracellular auxin compartment effects. The paper nevertheless keeps its main claim anchored in root auxin because that slice most directly exercises the multi-scale causal core.

The next step is not to argue that gene-driven emergence can run, but to use the validated core prospectively: test predicted mutants *in vivo* and scale execution by vectorizing the scheduler so larger cell populations can be simulated without losing benchmark discipline, debugger visibility, or causal interpretability. *The genetic program is the executable specification, and phenotype is the output of its execution within explicitly validated domains*.

## Notes

### Competing Interest Statement

The authors have declared no competing interest.

